# Fungal community turnover along steep environmental gradients from geothermal soils in Yellowstone National Park

**DOI:** 10.1101/2021.04.06.438641

**Authors:** Anna L. Bazzicalupo, Sonya Erlandson, Margaret Branine, Megan Ratz, Lauren Ruffing, Nhu H. Nguyen, Sara Branco

## Abstract

Geothermal soils offer unique insight into the way extreme environmental factors shape communities of organisms. However, little is known about the fungi growing in these environments and in particular how localized steep abiotic gradients affect fungal diversity. We used metabarcoding to characterize soil fungi surrounding a hot spring-fed thermal creek with water up to 84 °C and pH 10 in Yellowstone National Park. We found a significant association between fungal communities and soil variable principal components, and we identify the key trends in co-varying soil variables that explain the variation in fungal community. Saprotrophic and ectomycorrhizal fungi community profiles followed, and were significantly associated with, different soil variable principal components, highlighting potential differences in the factors that structure these different fungal trophic guilds. In addition, *in vitro* growth experiments in four target fungal species revealed a wide range of tolerances to pH levels but not to heat. Overall, our results documenting turnover in fungal species within a few hundred meters suggest many co-varying environmental factors structure the diverse fungal communities found in the soils of Yellowstone National Park.

## Introduction

Soil abiotic factors are known to strongly influence global fungal diversity [1], with soil moisture and chemistry as key drivers of large-scale fungal richness and composition [2]. However, little is known about whether these same drivers function to structure fungal communities at small scales, especially in environments with dramatic small-scale soil abiotic gradients. Geothermal areas are ideal systems to investigate such effects. Thermal sites are geologic units characterized by hot geothermal water, steam, and gases such as carbon dioxide, sulfur dioxide, and hydrogen chloride [3]. They are also notable for increased ambient soil temperature and landscape level thermal features such as hot springs and fumaroles. Thermal areas are by definition hot but can vary greatly in temperature ranging from mildly hot (∼30°C) to boiling point. In addition, geothermal water can cover the range of the pH scale [3]. Thermal soils can therefore display strong edaphic gradients within localized areas, with high variation in soil chemistry, moisture, and temperature.

Thermal areas and hot springs in particular are known for hosting diverse and specialized bacterial and archaeal communities, composed of thermophilic lineages, including novel ones, that evolved to tolerate extreme environmental conditions [4-7]. However, much less is known about microbial eukaryotic communities from thermal areas [8]. For the most part, members of Eukarya lack thermally stable membranes and are much less resilient to high temperatures compared to Archaea and Bacteria [9]. While the vast majority of Fungi fit this pattern, a very small number of thermophilic fungi have been documented. Thermophily evolved independently multiple times across the fungal tree of life [10] and a few species are able to withstand temperatures up to ∼60 °C [11]. Thermophilic fungi have been found in a wide variety of habitats responding to a range of different environmental pressures and can also occur outside of thermal areas [12]. Additionally, fungi are well known to withstand other extreme abiotic factors including broad pH ranges [13]. Several fungal species are able to grow within 5-9 pH unit differences even when originating from non-extreme habitats [14-16], suggesting that at the local scale, pH does not contribute strongly to structuring fungal communities. Thermophily in fungi is rare, so it is likely that in thermal areas, soil temperature may be the main factor driving fungal richness and composition, while other parameters such as pH are expected to be less relevant for structuring fungal diversity.

Little is known about the soil fungi from Yellowstone National Park (YNP)’s thermal habitats that are famous for extremophile research. Available studies have been either based on fungal culturing from soil (known to detect only a small portion of fungal diversity) or used low-resolution molecular approaches that limit the ability to fully detect fungal species diversity [17-20]. Only one very recent metagenomic study describes the mycobiome diversity of two hot springs in North Sikkim in India where they found the environments dominated by genera of Ascomycota and detected environmental yeast species [21]. Here we report on the effects of steep, localized soil abiotic gradients on fungi from Rabbit Creek, an alkaline thermal area in YNP. We expected geothermal water to form strong abiotic gradients forming in surrounding soils, directly impacting soil fungal diversity. Specifically, we hypothesized that more extreme soil conditions (i.e. higher temperature and pH) will host different and species-depauperate fungal communities relative to less extreme conditions. Given the low fungal tolerance to high temperatures, we predicted temperature to be the main factor affecting fungal diversity in this site. We used amplicon sequencing of the fungal ITS rRNA gene to characterize the fungal communities in soils across a gradient surrounding a hot spring-fed thermal creek with water up to ∼85 °C and pH ∼10. We found soil temperature was much lower than that of nearby boiling thermal water and covaried with moisture and pH. We found a significant association between fungal community and environmental soil variables. Although no single variable was a determining factor driving fungal community composition, soil variables were highly colinear allowing us to identify major soil chemistry trends that explained turnover in fungal communities. However, we found saprotrophic fungi and mycorrhizal (tree-associated) fungi were significantly associated with different sets of soil variables. In addition, we conducted *in vitro* growth assays testing for temperature and pH tolerance in four target fungal species (three saprobes and one mycorrhizal species) from Rabbit Creek. Although none of our target fungi were found to be thermophilic, they strongly differed in the ability to tolerate high or low soil pH.

## Methods

### Sampling site and sample processing

We sampled soils around Rabbit Creek, located in the Yellowstone National Park Lower Geyser Basin, an area characterized by numerous thermal features. Rabbit Creek flows directly out of a set of hot springs (Table S1, Figure S1). Water temperatures in Rabbit Creek start at approximately 84 °C at the main hot spring source and cool to 30 °C downstream. The water stays consistently alkaline at approximately pH 10. Site vegetation consisted of *Pinus contorta* forest with herbaceous plants including the heat tolerant hot springs panic grass, *Dichanthelium lanuginosum* (see Stout and Al□Niemi [22] for a complete geothermal plant survey of Yellowstone National Park).

In September 2018, we collected a total of 70 soil cores (2.5 × 10 cm) along 14 transects in the Rabbit Creek area. For each 20 m transect we sampled a total of five cores with one core every 5 m. We sampled eight transects perpendicular to Rabbit Creek (T2-T8) that were 150 m apart along the creek. We also collected six transects perpendicular to three nearby hot springs (two transects per hot spring, T1, T9-T13) and an additional transect (T14) away from surface water and in the pine forest (Table S1).

We measured the soil temperature 10 cm deep next to each soil core and homogenized the soil in each core before flash freezing and storing approximately 2 g of soil. We saved the remaining soil for chemical analyses. All cores were processed within 24 h of collection. Soil chemistry and moisture analyses were conducted at the Environmental Analytical Laboratory (Land Resources and Environmental Sciences, Montana State University). See Tables S1 and S2 for complete list of measurements.

### DNA extraction and library preparation for sequencing

We extracted genomic DNA from soil with the QIAGEN DNA Power Soil Pro kit using 0.5 g of soil instead of the recommended 0.25 g. For library preparation, we amplified the ITS1 region as it detects more taxa than ITS2 and recovers the same community patterns [23]. We used indexed primers ITS1F [24] and ITS2 [25] in the first round of PCR, followed by a second PCR to attach the 8 bp barcodes and Illumina adapters, using Hi-Fidelity Phusion DNA Polymerase for both reactions. Between each amplification step the product was cleaned from reagents and primers using the NGS SPRI Bead clean-up kit (ABM). Negative PCR controls were sequenced along with the experimental samples [26]. We included a synthetic mock community for ITS1 as a positive control to recover tag-switching among samples [27]. Amplicons of equimolar concentrations were pooled from each of 71 samples. The libraries were sent for paired 250-nucleotide reads on the Illumina MiSeq sequencing platform at the UC Davis DNA Sequencing Facility (Davis, California).

### Sequence processing

Raw sequence reads were demultiplexed at the UC Davis DNA Sequencing Facility (Davis, California). We used the QIIME2 software package [28], and a modified QIIME2 pipeline for fungal ITS by Nguyen [29] with some additional modifications. We quality filtered the demultiplexed raw sequences (q > 30) setting the truncation of sequences 240 bp for forward and 200 bp for reverse sequences, and denoised the raw reads with DADA2 [30]. After quality filtering, we *de novo* clustered sequences into operational taxonomic units (OTU) with VSEARCH at a 97% sequence similarity followed by a re-clustering step with UCLUST at 97% to ensure the best recovery of our OTUs according to our input mock community [26]. We then excluded all sequences that were ≤ 85% coverage in the BLAST search since this primer set amplifies sequences that only partially match to a small part of the ITS gene. We used the UNITE database v8, [31, 32] for chimera checking, OTU clustering, and assigning taxonomy.

Initial ordination analyses of the OTU community matrix led us to exclude 30 samples as they clustered due to partial, very short sequences spanning only ∼20 nucleotides at the beginning of the gene making them unidentifiable. In short, an initial non-metric dimensional scaling (NMDS) ordination including all remaining samples clustered in three groups (Figure S2). A species indicator analysis using *indicspecies* package (ver. 1.7.8) [33] in R with 9999 permutations showed the group labelled ‘3’ (Figure S2) clustered due to shared unidentifiable sequences and in fact, the ITS1 sequences for the OTUs clustered in this group because they could not be identified. We confirmed this by performing additional BLAST searches. We therefore excluded cores S5, S6, S7, S8, S9, S60, S63, S64, S67, and S69 from subsequent analyses. These cores were sampled near the three pools and the forest transect away from surface water (Table S1 and S2). After filtering, we used 40 cores to perform all further analyses. We produced a species accumulation curve using the *specaccum* function estimating the mean curve and its standard deviation from 100 random permutations of the species accumulation data. All scripts and data tables are available online at https://github.com/abazzical/YellowstoneFungi.

### Soil Chemistry and Fungal Community Analysis

We performed Principal Components Analysis (PCA) to assess chemistry similarities across soil cores. To investigate how the fungal community correlated with soil variables we performed a Redundancy analysis (RDA) on the OTU table and the first two PCs of the PCA using the vegan package [34, 35]. For the RDA, we transformed the OTU table by a centered log ratio (CLR).

We first added 1 to the OTU table to make the log transformation possible, then calculated the CLR transform with the *compositions* R package [36]. We categorized fungi into different trophic guilds using FUNGuild [37]. We further used the ectomycorrhizal (EM) OTUs and saprotrophic OTUs for additional RDAs to untangle which variables are more important for each fungal guild. To test for significant differences in the RDA we performed an ANOVA and controlled for transects with 1000 permutations within blocks.

To summarize species richness, we calculated Shannon Index in the *vegan* R package [34] and we report OTU numbers by rarefying the raw OTU table to a sampling depth of 2831. We tested if the species richness changed with distance from the edge of Rabbit Creek with a linear regression using the core number as the predictive variable.

### Fungal culture isolation and growth assays

We obtained pure cultures for four species growing at the Rabbit Creek site along the transect sampling sites. We cultured the saprotrophic *Agaricus campestris* and ectomycorrhizal *Pisolithus tinctorius* from fruitbodies on Potato Dextrose Agar (PDA) and Modified Melin-Norkans (MMN, [38]) media respectively. We also isolated cultures of potential plant pathogens *Fusarium oxysporum* and *F. avenae* from soil by plating serial soil dilutions in PDA. For all four species, we measured mycelial radial expansion at 30, 35, and 40°C and pH 4, 7, and 9 separately. Each culture was grown in triplicate and measured at 2-4 day intervals. *Fusarium* and *Agaricus* cultures were grown on PDA and *Pisolithus* was grown on MMN media, more suitable for mycorrhizal species. The growth in pH was calculated as the average of four radius measurements per colony for each time point. The temperature measurements were made by calculating the area of the fungal colony on the agar plate by importing the drawn outline of the fungus into ImageJ [39, 40].

We obtained culture identifications by sequencing the ITS rRNA gene. In short, we extracted genomic DNA using the Extract-N-Amp Sigma kit and amplified the ITS region with primers ITS1F [24] and ITS4 [25]. The PCR product was Sanger sequenced at GenScript and the sequences were checked and trimmed for low quality bases in the software FinchTV (http://www.geospiza.com/finchtv/). To identify the fungi, we used the BLAST tool in the NCBI database. Sequences are deposited in GenBank (MW471687 - MW472279).

## Results

### Soil abiotic variation

Soils in the Rabbit Creek area showed wide variation in temperature, moisture content, and chemistry (Fig. 1, S3). Despite the creek’s very high temperature water sources (84°C), soil temperature was relatively low and varied between 10 °C and 31°C. Soil pH and moisture content varied broadly, with pH ranging from 4 to 10 and moisture from 10% to 100%. pH and moisture contributed more than expected to PCA axis 1 and 2, respectively. Soil chemical parameters varied considerably (Table S1 and S2). Lead (Pb), iron (Fe), total carbon (TC), manganese (Mn), cobalt (Co), cadmium (Cd) were inversely proportional to alkaline values of pH and were contributing more than expected to PC1. Phosphorous (PO_4_), total nitrogen (TN), moisture content, copper (Cu), total carbon (TC), and zinc (Zn) were orthogonal to the first set of parameters and were contributing more than expected to PC2.

**Figure 1.**
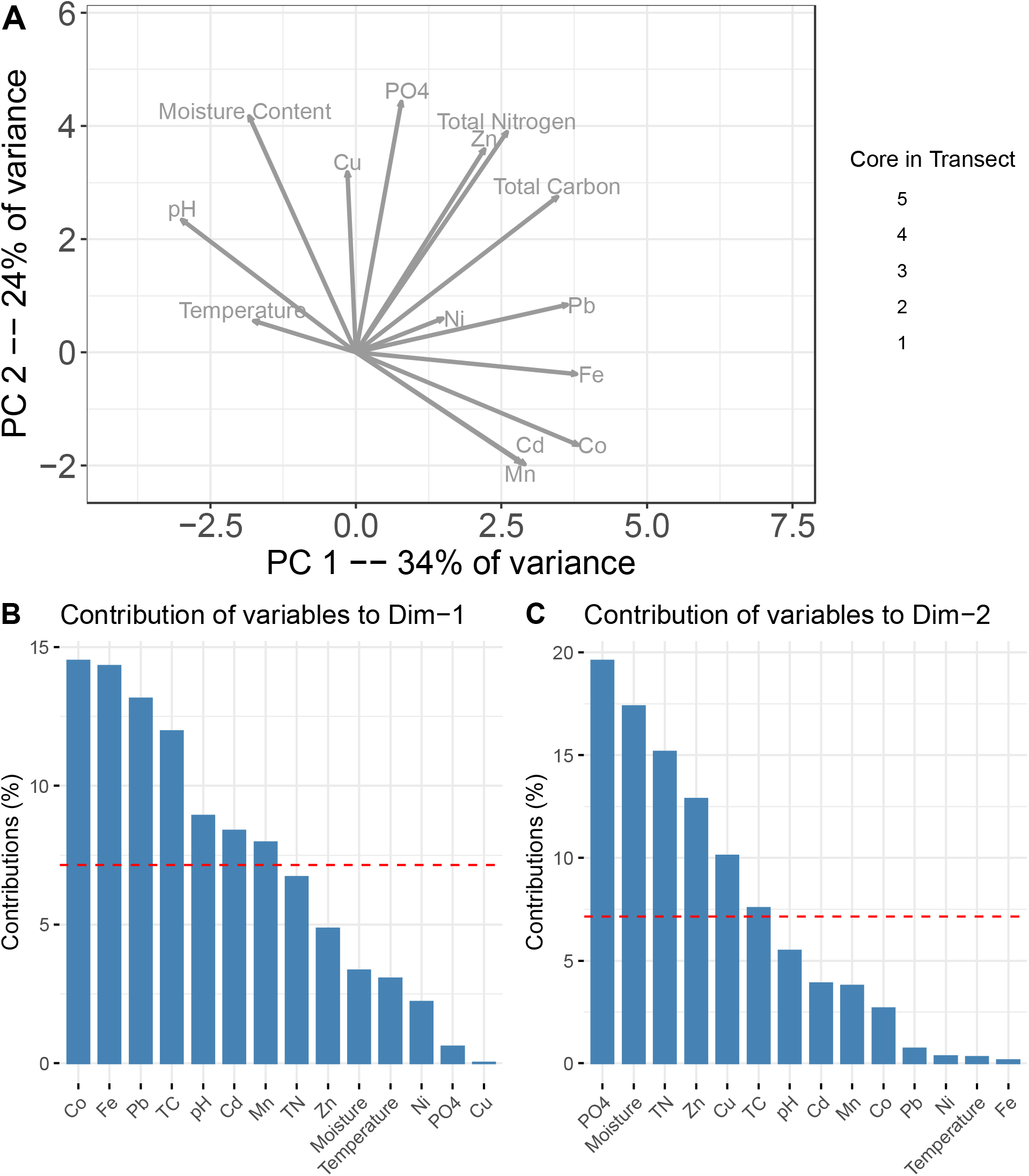
Rabbit Creek soil chemistry variation. (A) Principal components analysis on abiotic parameters measured across soil samples. Colors reflect the soil core number in each transect, core 1 is the closest to Rabbit Creek and core 5 is the furthest away (20 m from the shore). (B) Contribution of variables for PC1 and (C) PC2. Variables with contribution proportion above the red line contribute more than expected to a principal component.

### Overall soil fungal community composition

We found a total of 593 fungal OTUs at Rabbit Creek and were able to obtain taxonomic identifications at a lower taxonomic level than “fungi” for close to 65% of them. This thermal site was heavily dominated by members of Ascomycota (38.4%) and Basidiomycota (18.7%) but also included Chytridiomycota, Rozellomycota, Mucoromycota, Glomeromycotina, Kickxellomycota, Mortierellomycota, and Calcarisporiellomycota. We detected a few fungi known for thriving at high temperatures (>30 °C), but not resilient enough to be considered heat tolerant fungi (with the ability to grow at 40 °C [41]). These included *Exophiala opportunistica*, a black yeast that also withstands high moisture and alkalinity and is commonly found in dishwashers [42, 43], *Waitea circinata*, a grass pathogen with optimal growth at 25-30 °C [44], *Umbelopsis vinacea*, a soil fungus that remains active up to 37 °C [45], *Gaertneriomyces semiglobifer*, a chytrid that also growth up to 37 °C [46], and *Fusarium kerasplasticarum*, an opportunistic animal pathogen that can also grow up to 37 °C. In addition, we found one unidentified species of *Talaromyces*, a genus that includes thermophilic and thermotolerant species [47]. Notably, we recovered a number of cold-adapted fungi in our survey, including *Extremus aquaticus, Elasticomyces elasticus, Endophoma alongata, Alternaria chlamydosporigena, Mrakiella aquatica, Mrakia frigida, Tausonia pamirica, T. pulullans, Naganishia adeliensis*, and *Udeniomyces megalosporus*. Many of these species occur in Polar regions and have documented optimal growth temperatures <20 °C [48-53].

Our species accumulation curve based on samples does not plateau (Figure S4), suggesting that community diversity is not saturated with our sampling.

### Redundancy analysis

We performed an RDA between the OTU matrix and the first two PCs of the environmental variables PCA and we found fungal communities were significantly different and correlated with the first two PCs of the environmental variables PCA (Figure 2, Table 1). The adjusted R^2^ for this model was 0.5 (Table 1).

**Figure 2.**
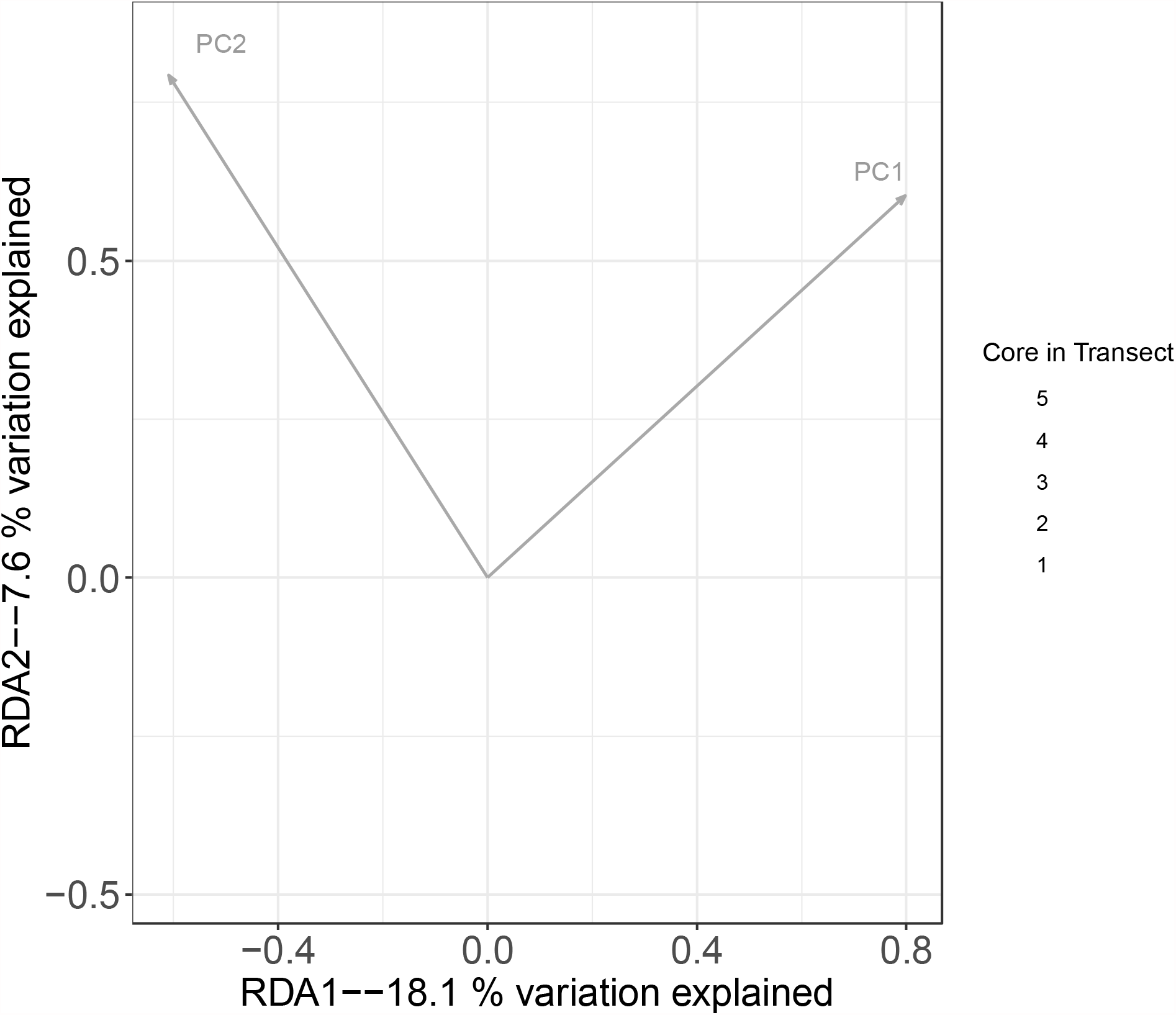
Redundancy analysis of the fungal community and first two PCs of the soil environmental variable PCA. Both PCs contributed to RDA axes and significantly explained community composition. The variance explained by each RDA axis is for fitted (constrained) values. Colors reflect the soil core number in each transect, core 1 is the closest to Rabbit Creek and core 5 is the furthest away (20 m).

**Table 1.** Summary of RDA full model analysis of variances (1000 permutations), proportion of fitted variance explained by RDA axes 1 and 2 for each model, and analysis of variance of individual predictors (soil principal components) in each RDA model.

| **RDAs summary statistics** |  |  |  |  |  |  |
| --- | --- | --- | --- | --- | --- | --- |
|  | All OTUs | residuals | Saprotrophic | residuals | EM | residuals |
| Adjusted R2 | 0.05 | -- | 0.094 | -- | 0.057 | -- |
| DF | 2 | 45 | 2 | 35 | 2 | 35 |
| Variance | 25.70 | 269.26 | 8.70 | 51.76 | 5.10 | 41.80 |
| F | 2.15 | -- | 2.94 | -- | 2.14 | -- |
| Pf(>F) | **0.000999 \*\*\*** | -- | **0.02597 \*** | -- | **0.004995 \*\*** | -- |
| **RDA1 and RDA2 contributions** |  |  |  |  |  |  |
|  | All OTUs |  | Saprotrophic |  | EM |  |
|  | RDA1 | RDA2 | RDA1 | RDA2 | RDA1 | RDA2 |
| Eigenvalue | 18.12 | 7.57 | 6.18 | 2.52 | 3.39 | 1.72 |
| Proportion Explained | 0.71 | 0.29 | 0.71 | 0.29 | 0.66 | 0.34 |
| Cumulative Proportion | 0.71 | 1 | 0.71 | 1 | 0.66 | 1 |
| **Individual significance of Principal Components** |  |  |  |  |  |  |
|  | All OTUs |  | Saprotrophic |  | EM |  |
|  | PC1 | PC2 | PC1 | PC2 | PC1 | PC2 |
| DF | 1 | 1 | 1 | 1 | 1 | 1 |
| Variance | 16.00 | 9.97 | 5.54 | 3.29 | 3.39 | 1.73 |
| F | 2.67 | 1.67 | 3.75 | 2.22 | 2.83 | 1.45 |
| Pf(>F) | **0.000999 \*\*\*** | **0.023976 \*** | 0.07 | **0.000999 \*** | **0.004995 \*** | 0.63 |

Two hundred and thirty-four (out of 593) OTUs were classified into trophic guilds with FUNGuild. Twenty two percent (52) OTUs were mycorrhizal (including 16.7% (39) EM fungi), 34.6% (81) saprotrophic, 9.8% (23) plant pathogens, 15% (35) animal pathogens, and 6.8% (16) dung-associated fungi (Table S3). We performed RDAs on OTU tables composed only of saprotrophic and EM fungal taxa (Figure 3). Both guilds were found to be significantly associated with soil variables PCs (Table 1), however, saprotrophic fungi were only significantly associated with PC2 while EM fungi were significantly associated only with PC1. The total variance explained (adj. R^2^) in the RDA was 9.4% for saprotrophic fungi and 5.7% for EM fungi (Table 1).

**Figure 3.**
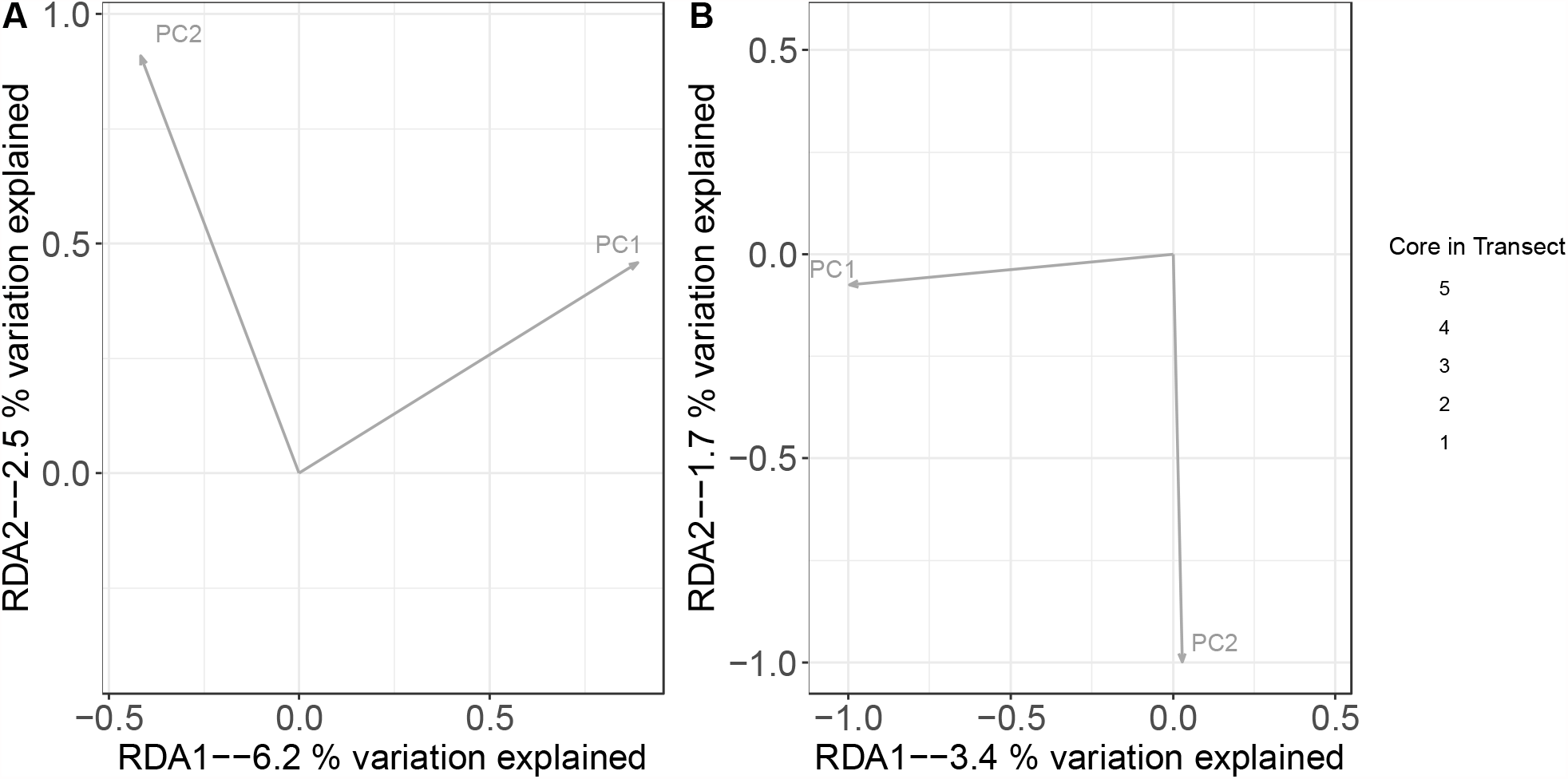
Ectomycorrhizal (EM) and saprotrophic fungal community redundancy analyses. (A) Saprotrophic fungi are significantly associated with PC2. (B) EM fungi are significantly associated with PC1. Variables contributing more than expected to PC1 were: pH, Lead (Pb), iron (Fe), total carbon (TN), manganese (Mn), cobalt (Co), and cadmium (Cd). Variables contributing more than expected to PC2 were: Phosphorous (PO_4_), total nitrogen (TN), moisture content, copper (Cu), total carbon (TC), and zinc (Zn).

### Species richness measures

We report OTU numbers and Shannon Index measurements per core. The cores of each transect represent a soil environmental gradient moving away from the creek shore that is supported by the soil PCA. We tested if the distance from the creek edge affected the fungal species richness. We performed linear regressions for the Shannon Index and the OTU numbers against the core number which reflects the distance from the creek edge. However, we do not find a changing pattern of species richness away from the creek shore suggesting a change in community turnover rather than change in number or richness of species (Figure 4). None of the linear models showed overall significance, however, we found two significant differences for a single predictive variable in two groups: in OTU numbers for core 2 when we tested all the OTUs (Estimate=15.929 Std. Error=6.332 t value=2.516 Pr(>|t|)=0.0169) and the saprotrophic OTUs (Estimate=6.119 Std. Error=2.099 t value=2.915 Pr(>|t|)=0.006344).

**Figure 4.**
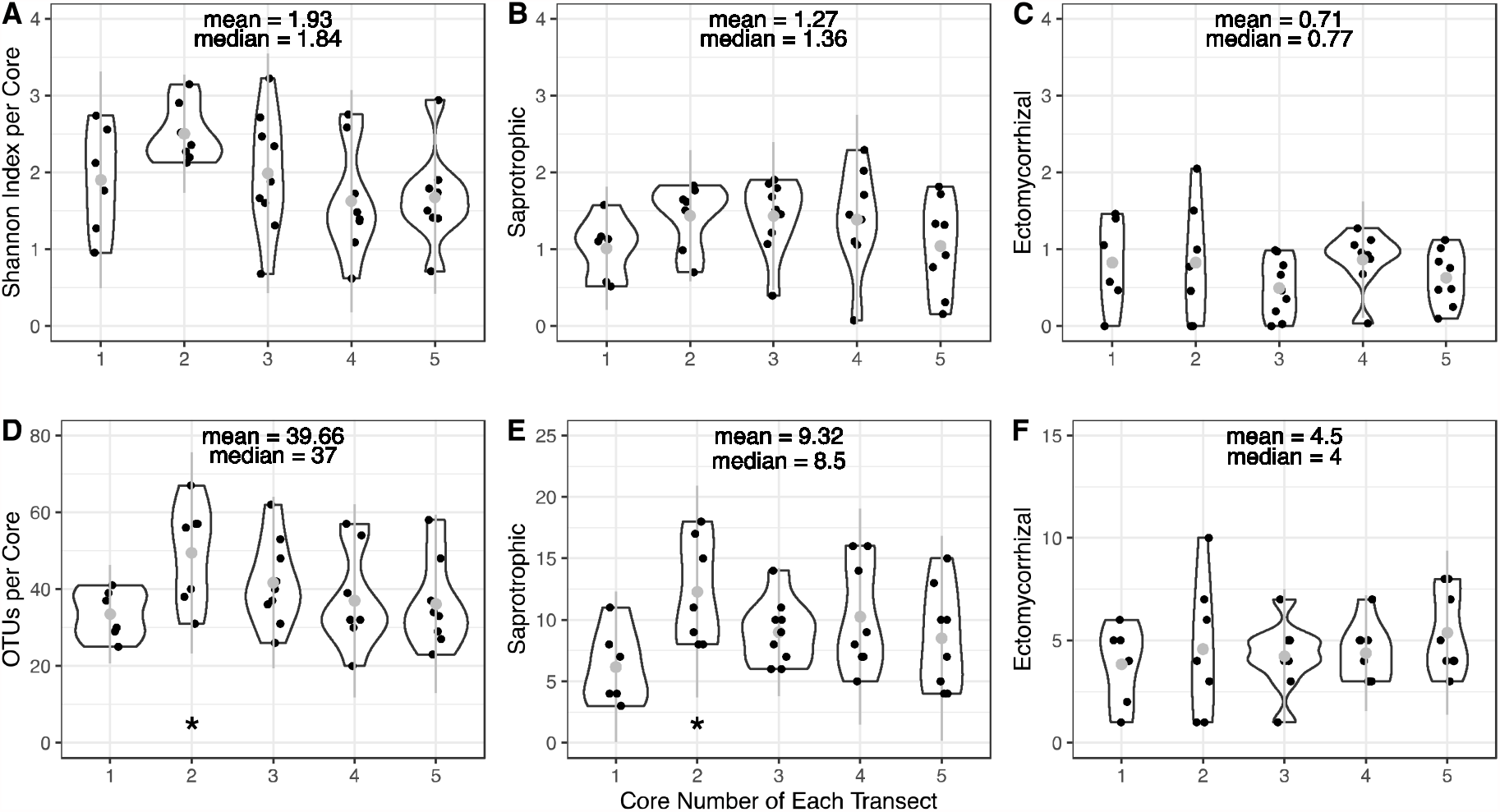
Shannon Index and OTU numbers by distance from the shore of Rabbit Creek. Core number in each transect correspond to core 1=closest to the water’s edge ∼0m, core 2=5m from the shore, core 3=10m, core 4=15m, core 5=20m. (A) Shannon Index of all fungal OTUs; (B) Shannon Index of saprotrophic fungal OTUs; (C) Shannon Index of EM fungal OTUs; (D) number of all fungal OTUs; (E) number of saprotrophic fungal OTUs; (E) number of EM fungal OTUs. Note the scale bar differences in D-F. Asterisks (^*^) indicate significance.

### Effect of temperature and pH on four fungi from Yellowstone National Park soils

The four fungal species tested for temperature and pH tolerance lacked thermal tolerance but had the ability to tolerate a wide range of pH (Figures 5 and 6). Notably, all four species performed well at pH 7. Both *Fusarium oxysporum* and *F. avenae* grew equally well at pH 9 and pH 7 showing less growth at pH 4. Both *Agaricus campestris* and *Pisolithus tinctorius* grew best at pH 7, however, *P. tinctorius* was the only fungus with no growth at pH 9 (Figure 5). All four species grew well at 30 °C but very poorly at 40 °C (Figure 6). *P. tinctorius* and *F. oxysporum* grew equally well at 30 °C and 35 °C, while *A. campestris* and *F. avenae* grew better at 30 °C than 35 °C. Unfortunately, we did not detect any of the taxa we cultured in our metabarcoding data. This is not very surprising given the species accumulation curves do not plateau (Figure S4) indicating our environmental sequencing only detected part of the fungi inhabiting the Rabbit Creek soil.

**Figure 5.**
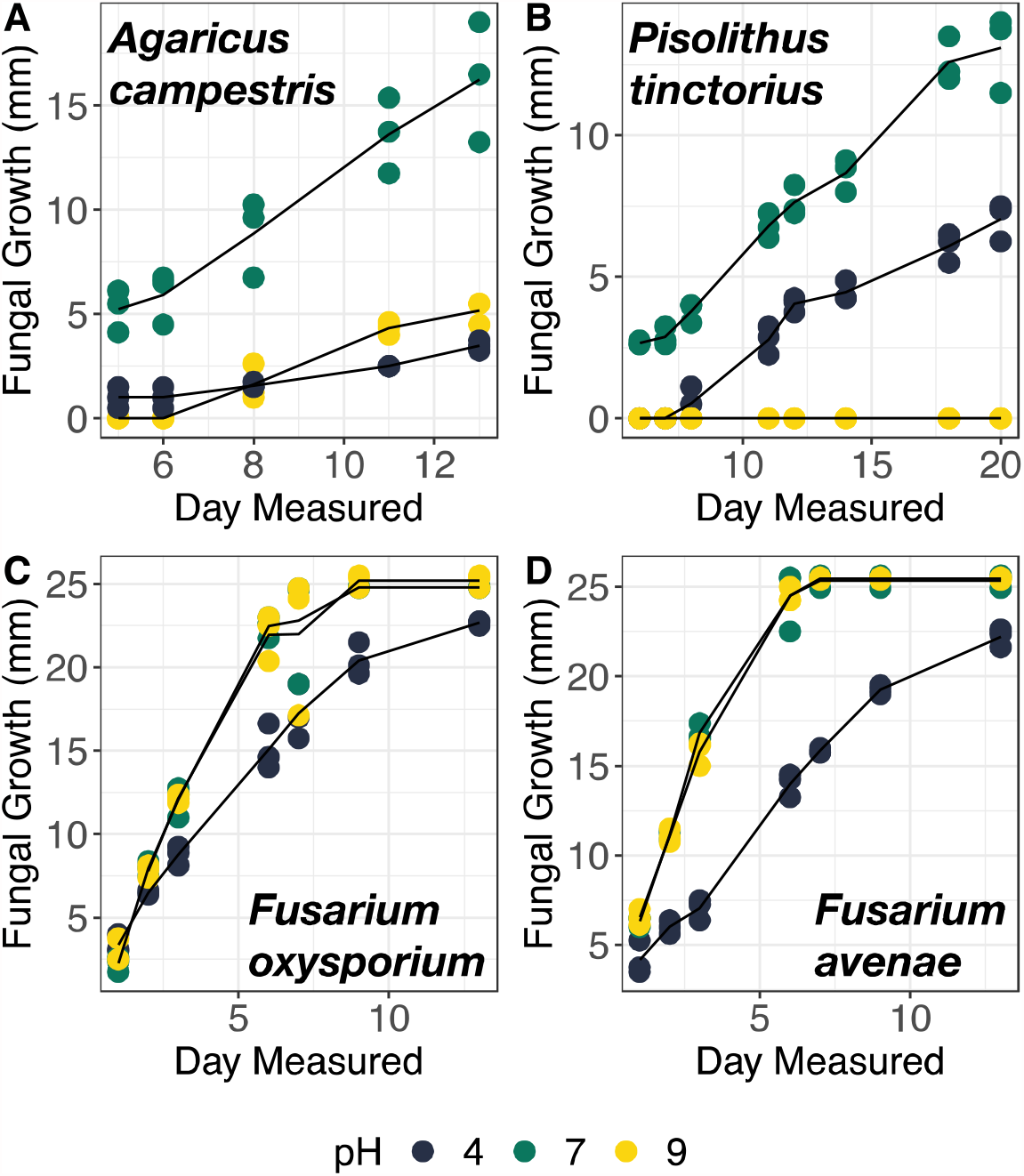
Mycelial growth (average of 4 radius measurement per fungal thallus) at pH 4 (dark blue), pH 7 (green), and pH 9 (yellow) in *Agaricus campestris* (A), *Pisolithus tinctorius* (B), *Fusarium oxysporum* (C), and *Fusarium avenae* (D).

**Figure 6.**
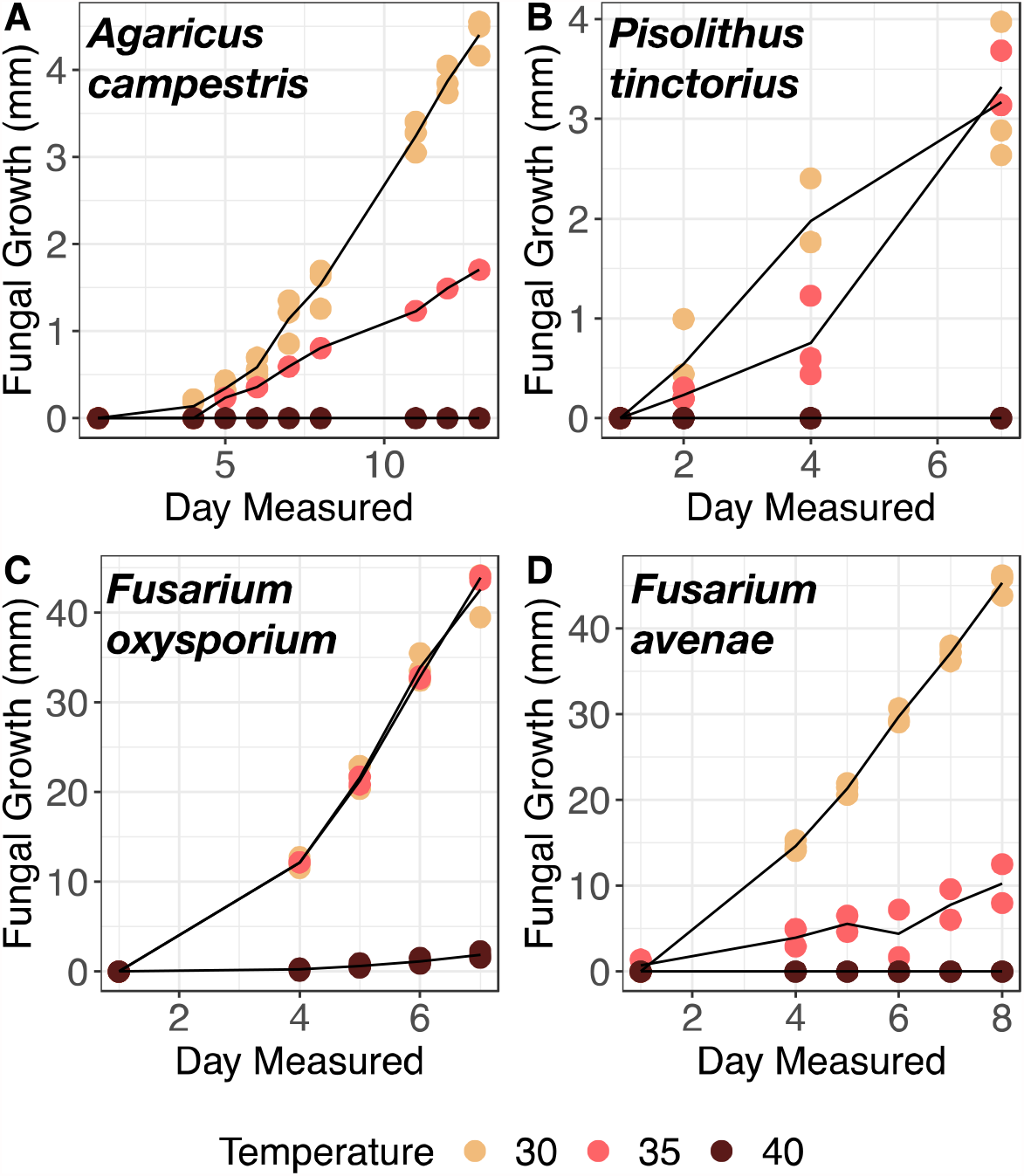
Mycelial growth (fungus growth area mm^3^) at 30°C (peach), 35°C (pink), and 40°C (burgundy) in *Agaricus campestris* (A), *Pisolithus tinctorius* (B), *Fusarium oxysporum* (C), and *Fusarium avenae* (D).

## Discussion

We investigated soil fungal communities from a geothermal area in Yellowstone National Park characterized by a steep abiotic gradient and found a significant association between fungal communities and soil environmental variables. We summarized the highly co-varied soil environmental data with PCA and the first two environmental variable PCs significantly explained variation for the whole fungal community. In addition, saprotrophic and EM fungi were significantly associated with different components of soil variables. It is known that global and continental patterns of fungal diversity are shaped by edaphic factors [1, 54]. We present evidence that the same co-varying edaphic factors that structure fungal diversity at the global scale also structure fungal communities at a scale of only a few hundred meters. In our study, these edaphic factors included gradients of co-varied pH, moisture, heavy metals, phosphate, total carbon and nitrogen. We also found no decline in number or richness measure for OTUs suggesting that species turnover rather than a reduction in diversity are responsible for the patterns we reported.

We were surprised by the absence of thermophilic fungi and the presence of cold-adapted lineages in Rabbit Creek. On the contrary, we detected several cold-adapted species previously recorded in the region. Specifically, we found the psychrophilic basidiomycete yeast *Tausonia pulullans* [55] and a few cold adapted Ascomycete yeasts identified in Vu *et al*. (2016) [56], including *Naganishia adeliensis* and *Udenomyces megalosporus* (=*U. pyricola* in Vu et al 2016). We also found the psychrophiles *Elasticomyces elasticus* and *Mrakia sp*., that had been detected in previous soil surveys in both West Yellowstone and Hyalite near Cooke City, Montana [57]. Additionally, our *Tausonia pamirica* OTU matched 100% to the sequence of the type specimen (Genbank # NR_154490), a known psychrotolerant yeast from Antarctica [58]. The occurrence of these species is most likely unrelated to the thermal nature of Rabbit Creek and rather may reflect the regional climate characterized by severe cold winters. We were also surprised that none of the fungi we isolated in culture were able to grow at temperatures higher than 35°C. Soil temperature is thought to be the most significant factor influencing bacterial and archaeal species composition in geothermally heated environments [22, 59, 60]. Thermophilic fungi are mostly known from studies on plant-associated fungi from geothermal environments [61] and surveys such as the YNP metagenome project [62]. There is also experimental evidence documenting previously unknown thermophilic and acidophilic fungal species from hot spring habitats [63].

However, the soil temperature of our sampled cores was relatively low, ranging from 10 °C to 31°C, which is well within the range of known temperature tolerance of most fungi including the detected psychrophilic species [55, 56] and the fungi tested in this study (Fig. 6). It is important to note that our findings do not imply the absence of heat-adapted fungi in Yellowstone, as other thermal areas characterized by warmer soils may well host thermophilic fungi of unknown taxonomic identity.

Conditions in highly acidic or alkaline environments often co-vary with other environmentally extreme variables such as elevated salt concentrations or temperatures[64], and in our case, soil moisture content. It is known that edaphic parameters structure communities of soil fungi, and variation in chemistry (including pH) affects fungal diversity on global, continental and local scales [1, 2]. Individual fungi appear to have a wider pH tolerance range compared to bacteria [65], however there are several examples of fungi (including *P. tinctorus*, which was studied here) being able to withstand both low and high pH levels [14]. Studies on acid- or alkaline-loving fungi have recorded new species and have expanded our knowledge of the possible habitat types for commonly found fungal species [13]. In addition, research on fungi associated with hot spring sediment across the globe has documented the phylum Ascomycota as dominant in these habitats. These included newly discovered species [63] and common members of widespread genera, such as *Aspergillus, Penicillium*, and *Cladosporium*, consistent with taxa found in hot spring environments in North Sikkim, India [21]. Mycorrhizal species belonging to lineage Ascomycota have also been detected soda lake habitats [66]. We found the Rabbit Creek soils with higher pH are inhabited by many members of the Ascomycota lineage, these include several alkalitolerant fungi such as *Acremonium* from the Emericellopsis clade [67].

We found differences on the effect of edaphic parameters across fungal guilds. EM fungal composition was significantly associated with the axis influenced by heavy metals, pH values, and carbon, while saprotrophic fungi were significantly associated with an axis influenced by phosphorous, nitrogen, moisture, carbon (to a lesser extent than the first axis), and copper and zinc. Additionally, the number of saprotrophic OTUs significantly changed with distance from the creek shore, with an uptick at an intermediate distance while there was no difference found in EM fungi. Interestingly, mycorrhizal fungi have been predicted to display more restricted ranges compared to saprotrophs [68], although global empirical data failed to recover this pattern [69]. This would lead to a prediction of edaphic factors affecting mycorrhizal fungi disproportionately, which is the opposite of what we found. Instead, our study suggests either that the ability to associate with a host allows mycorrhizal fungi to inhabit any micro-habitat where the host is present or that saprotrophic fungi might be more sensitive to environmental variables when resource competition and successional factors are removed.

In conclusion, we found statistical support suggesting soil variables structured fungal community composition at the scale of only a few hundred meters. We also found thermophilic fungi to be absent from the soils surrounding thermal water in Rabbit Creek. Further research on thermal areas characterized by extreme soil conditions is however needed improving our understanding of the effects of extreme edaphic factors on the ecology and evolution of fungi.

## Acknowledgements

We thank Maddie Trent, Rio Wofford, Colin Kennedy, Seamus Hoolahan, and Kathryn Gannon for assistance in the field and laboratory, and the Quandt lab for suggestions on the manuscript. Cathy Zabinski helped obtaining the collecting permit, Dan Colman assisted with sampling design, and Kabir Peay provided raw data from another study that allowed fungal community comparisons. In addition, we thank three anonymous reviewers for crucial input on previous versions of this study. We also thank Yellowstone National Park for allowing the collection of samples in Rabbit Creek (YELL-2018-SCI8062).

## Declarations

The authors declare no funding.

The authors have no conflicts of interest to declare.

The research did not require ethical approval, no consent to participate or publication was needed.

All sequences are in GenBank and will be made public after publication: MW471687 - MW472279.

Code for analyses is available at: https://github.com/abazzical/YellowstoneFungi

All authors have approved this draft of the paper.

