## Supplement 1 for "Fungal community turnover along steep environmental gradients from geothermal soils in Yellowstone National Park"

**Figures**

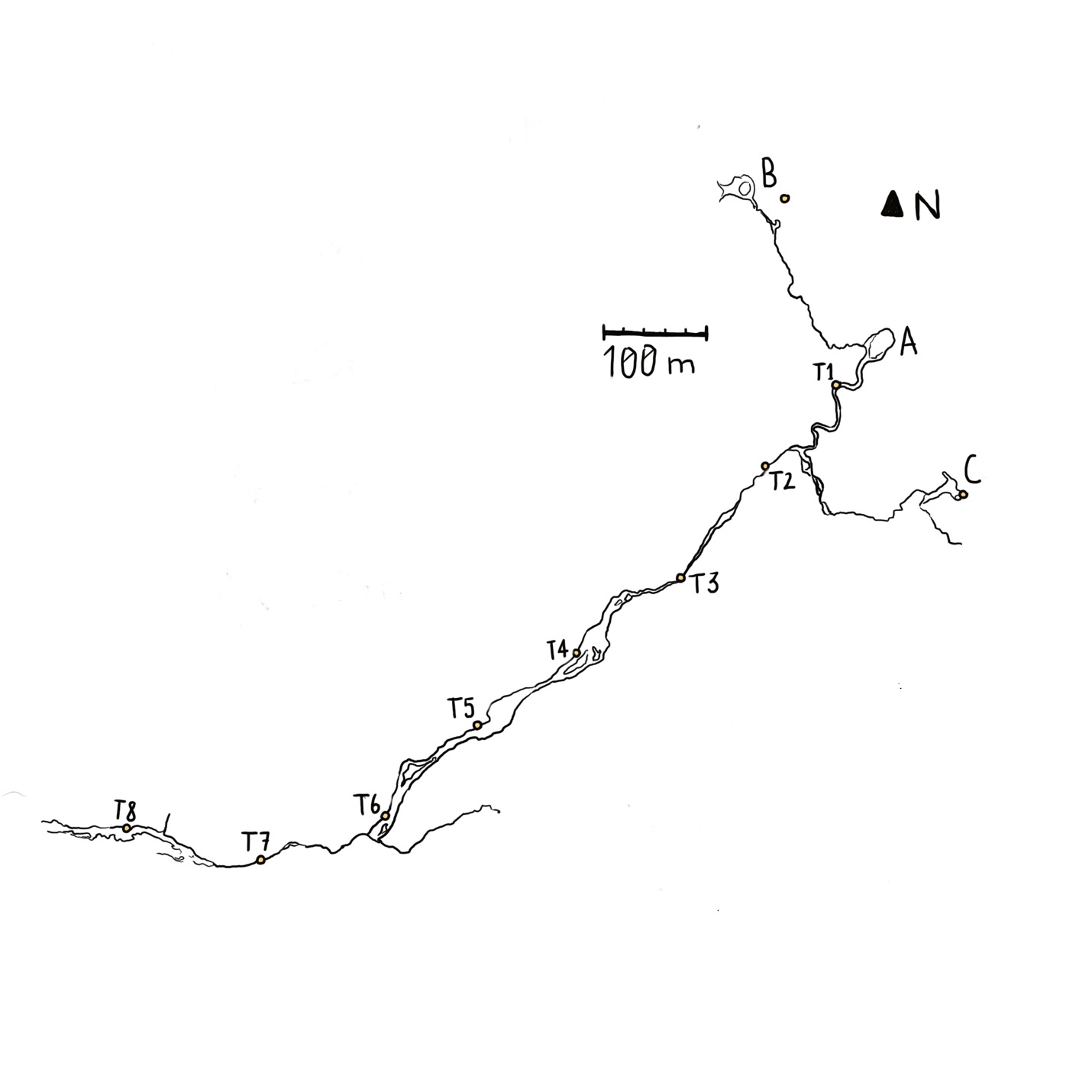

**Figure S1** Rabbit Creek sampling site. Transects (T) are marked at the start of each transect for the three pools (A, B, and C) and along the creek (N 44 30 W 110 49).

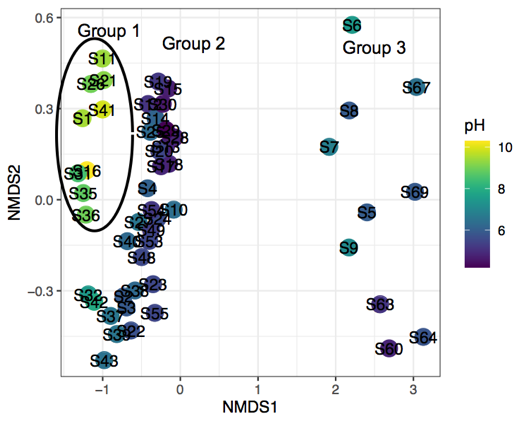

**Figure S2** NMDS including all cores forms three distinct clusters. The left-most cluster (including cores S5-9, S60, S63, S64, S67, S69) did not show any distinction based on environmental variables. An indicator species analysis found that 26 indicator species were all ‘unidentified’ taxa based on ITS1, and on further analysis of the sequences, we found them to be too short to be assigned to any particular taxon. Based on this result we cannot exclude that these cores were grouped in the NMDS based on incomplete data, we therefore excluded them from the following analyses.

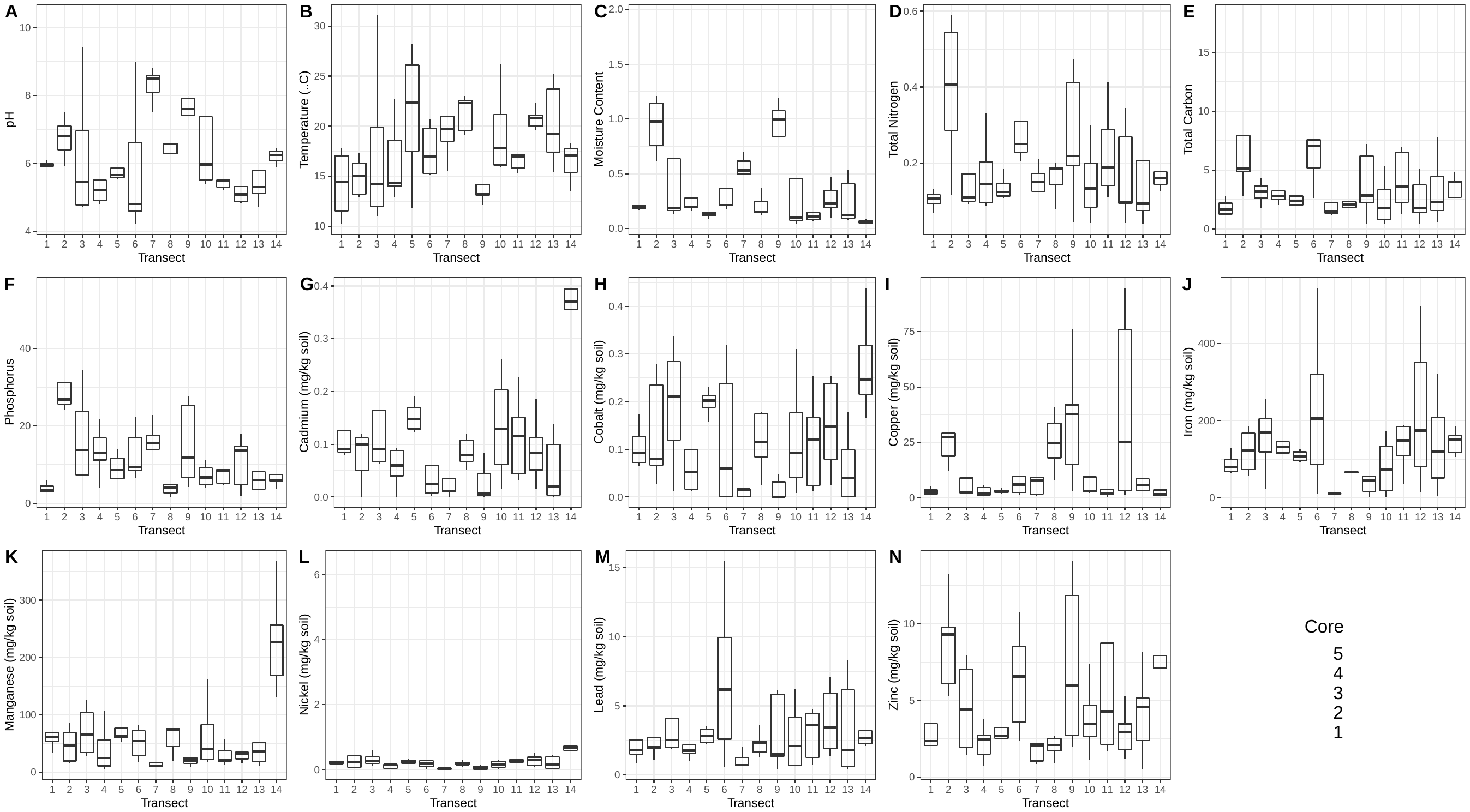

**Figure S3** Distribution of environmental factor measurements of soil cores.

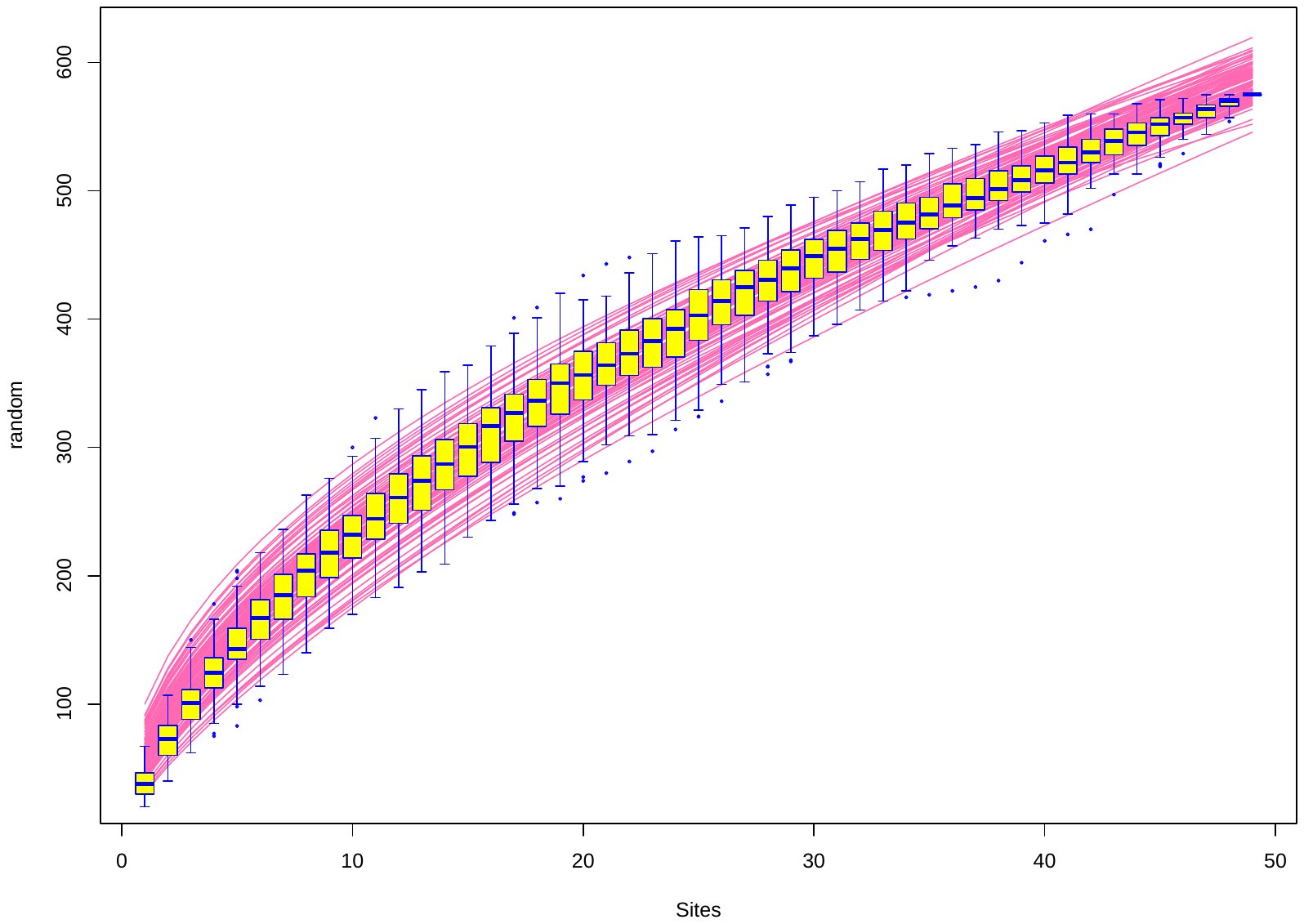

**Figure S4** Species accumulation curve of fungal soil communities collected in Rabbit Creek in Yellowstone National Park. Error bars are based on 100 random permutations of the data.

**Tables**

**TABLE S1** Soil core sample information for temperature, transect number, environment type (pool, transect from creek, forest), cores Filtered based on sequence quality used in NMDS: starred* cores are ones from Group 3 and therefore excluded in the final NMDS, number of OTUs per core and latitude and longitude coordinates.

| **ID** | **Temperature** | **Transect** | **Env. Type** | **Filtered** | **# OTUs** | **Lat** | **Long** |
| --- | --- | --- | --- | --- | --- | --- | --- |
| S1 | 17.1 | 1 | Pool A | S1 | 29 | 44 31.256 | 110 48.738 |
| S2 | 17.8 | 1 | Pool A | S2 | 56 | 44 31.257 | 110 48.740 |
| S3 | 16.8 | 1 | Pool A | S3 | 36 | 44 31.260 | 110 48.742 |
| S4 | 12 | 1 | Pool A | S4 | 32 | 44 31.262 | 110 48.745 |
| S5 | 10.2 | 1 | Pool A | S5* | 39 | 44 31.265 | 110 48.747 |
| S6 | 16.3 | 2 | Creek | S6* | 23 | 44 31.211 | 110 48.793 |
| S7 | 12.9 | 2 | Creek | S7* | 50 | 44 31.213 | 110 48.795 |
| S8 | 13.2 | 2 | Creek | S8* | 31 | 44 31.216 | 110 48.797 |
| S9 | 17.3 | 2 | Creek | S9* | 44 | 44 31.218 | 110 48.798 |
| S10 | 15 | 2 | Creek | S10 | 58 | 44 31.220 | 110 48.800 |
| S11 | 31.1 | 3 | Creek | S11 | 25 | 44 31.150 | 110 48.858 |
| S12 | 16.7 | 3 | Creek | S12 | 29 | 44 31.152 | 110 48.861 |
| S13 | 16.2 | 3 | Creek | S13 | 31 | 44 31.153 | 110 48.863 |
| S14 | 12.3 | 3 | Creek | S14 | 32 | 44 31.154 | 110 48.867 |
| S15 | 11 | 3 | Creek | S15 | 23 | 44 31.155 | 110 48.871 |
| S16 | 22.7 | 4 | Creek | S16 | 41 | 44 31.109 | 110 48.939 |
| S17 | 18.6 | 4 | Creek | S17 | 31 | 44 31.108 | 110 48.943 |
| S18 | 14 | 4 | Creek | S18 | 40 | 44 31.112 | 110 48.945 |
| S19 | 14.3 | 4 | Creek | S19 | 20 | 44 31.113 | 110 48.948 |
| S20 | 12.9 | 4 | Creek | S20 | 33 | 44 31.116 | 110 48.953 |
| S21 | 21.2 | 5 | Creek | S21 | 30 | 44 31.069 | 110 49.018 |
| S22 | 28.2 | 5 | Creek | S22 | 57 | 44 31.072 | 110 49.020 |
| S23 | 19.4 | 5 | Creek | S23 | 42 | 44 31.073 | 110 49.022 |
| S24 | 25.4 | 5 | Creek | S24 | 39 | 44 31.075 | 110 49.024 |
| S25 | 11.8 | 5 | Creek | S25 | 37 | 44 31.077 | 110 49.026 |
| S26 | 20.7 | 6 | Creek | S26 | 30 | 44 31 018 | 110 49.088 |
| S27 | 19.8 | 6 | Creek | S27 | 57 | 44 31.018 | 110 49.092 |
| S28 | 17 | 6 | Creek | S28 | 26 | 44 31.018 | 110 49.095 |
| S29 | 15.1 | 6 | Creek | S29 | 30 | 44 31.019 | 110 49.098 |
| S30 | 15.3 | 6 | Creek | S30 | 29 | 44 30.021 | 110 49.102 |
| S31 | 19.7 | 7 | Creek | S31 | 37 | 44 30.993 | 110 49.189 |
| S32 | 25.7 | 7 | Creek | S32 | 40 | 44 30.995 | 110 49.190 |
| S33 | 18.5 | 7 | Creek | - | - | 44 30.998 | 110 49.190 |
| S34 | 21 | 7 | Creek | - | - | 44 31.001 | 110 49.191 |
| S35 | 15.5 | 7 | Creek | S35 | 27 | 44 31.003 | 110 49.191 |
| S36 | 23 | 8 | Creek | S36 | 39 | 44 31.010 | 110 49.295 |
| S37 | 19.1 | 8 | Creek | S37 | 67 | 44 31.013 | 110 49.295 |
| S38 | 22.6 | 8 | Creek | S38 | 62 | 44 31.016 | 110 49.297 |
| S39 | 22.3 | 8 | Creek | S39 | 32 | 44 31.018 | 110 49.298 |
| S40 | 19.6 | 8 | Creek | S40 | 48 | 44 31.020 | 110 49.297 |
| S41 | 19.1 | 9 | Pool B | S41 | 29 | Not Recorded | Not Recorded |
| S42 | 14.2 | 9 | Pool B | S42 | 38 | Not Recorded | Not Recorded |
| S43 | 13.2 | 9 | Pool B | S43 | 37 | Not Recorded | Not Recorded |
| S44 | 13.1 | 9 | Pool B | - | - | Not Recorded | Not Recorded |
| S45 | 12.1 | 9 | Pool B | - | - | Not Recorded | Not Recorded |
| S46 | 26.2 | 10 | Pool B | - | - | Not Recorded | Not Recorded |
| S47 | 19.5 | 10 | Pool B | - | - | Not Recorded | Not Recorded |
| S48 | 16.2 | 10 | Pool B | S48 | 53 | Not Recorded | Not Recorded |
| S49 | 15.9 | 10 | Pool B | S49 | 54 | Not Recorded | Not Recorded |
| S50 | 17.8 | 10 | Pool B | - | - | Not Recorded | Not Recorded |
| S51 | 23.5 | 11 | Pool C | - | - | Not Recorded | Not Recorded |
| S52 | 15.8 | 11 | Pool C | - | - | Not Recorded | Not Recorded |
| S53 | 15.3 | 11 | Pool C | S53 | 48 | Not Recorded | Not Recorded |
| S54 | 17 | 11 | Pool C | S54 | 57 | Not Recorded | Not Recorded |
| S55 | 17.2 | 11 | Pool C | S55 | 34 | Not Recorded | Not Recorded |
| S56 | 20.8 | 12 | Pool C | - | - | Not Recorded | Not Recorded |
| S57 | 20 | 12 | Pool C | - | - | Not Recorded | Not Recorded |
| S58 | 22.3 | 12 | Pool C | - | - | Not Recorded | Not Recorded |
| S59 | 21.1 | 12 | Pool C | - | - | Not Recorded | Not Recorded |
| S60 | 19.6 | 12 | Pool C | S60* | 43 | Not Recorded | Not Recorded |
| S61 | 25.2 | 13 | Pool A | - | - | Not Recorded | Not Recorded |
| S62 | 23.7 | 13 | Pool A | - | - | Not Recorded | Not Recorded |
| S63 | 19.2 | 13 | Pool A | S63* | 45 | Not Recorded | Not Recorded |
| S64 | 17.4 | 13 | Pool A | S64* | 22 | Not Recorded | Not Recorded |
| S65 | 15.4 | 13 | Pool A | - | - | Not Recorded | Not Recorded |
| S66 | 13.5 | 14 | Forest | - | - | 44 30.459 | 110 49.936 |
| S67 | 18.3 | 14 | Forest | S67* | 20 | 44 30.452 | 110 49.937 |
| S68 | 17.8 | 14 | Forest | - | - | 44 30.452 | 110 49.941 |
| S69 | 15.4 | 14 | Forest | S69* | 26 | 44 30 453 | 110 49.945 |
| S70 | 17.1 | 14 | Forest | - | - | 44 30.454 | 110 49.948 |

**Table S2** Chemistry measurements for 70 soil cores from Rabbit Creek. The metal units are all mg metal per kg of soil (mg/kg soil). (Nickel measurement for sample 1.1 is missing. We used average of the nickel values for transect 1 to be able to run the PCoA).

| **ID** | **Sample** | **Moisture** | **pH** | **PO4** | **TN** | **TC** | **Cd** | **Co** | **Cu** | **Fe** | **Mn** | **Ni** | **Pb** | **Zn** |
| --- | --- | --- | --- | --- | --- | --- | --- | --- | --- | --- | --- | --- | --- | --- |
| S1 | 1.1 | 0.5 | 9.49 | 5.235 | 0.086 | 1.023 | 0.016 | 0.02 | 0.016 | 8.24 | 7.28 | - | 0.32 | 0.232 |
| S2 | 1.2 | 0.256 | 5.91 | 2.716 | 0.111 | 1.291 | 0.087 | 0.075 | 5.118 | 90.276 | 33.701 | 0.181 | 1.85 | 2.087 |
| S3 | 1.3 | 0.194 | 5.91 | 5.929 | 0.132 | 1.966 | 0.095 | 0.111 | 1.786 | 70.238 | 61.389 | 0.214 | 0.873 | 2.619 |
| S4 | 1.4 | 0.19 | 6.08 | 3.887 | 0.1 | 2.818 | 0.218 | 0.174 | 2.97 | 130.297 | 92.752 | 0.368 | 4.634 | 6.059 |
| S5 | 1.5 | 0.171 | 5.96 | 2.98 | 0.067 | 1.129 | 0.08 | 0.064 | 1.32 | 64.36 | 60.28 | 0.18 | 1.72 | 2.04 |
| S6 | 2.1 | 1.143 | 7.5 | 31.192 | 0.406 | 5.094 | 0 | 0.027 | 18.784 | 58.243 | 15.946 | 0.041 | 1.054 | 6.081 |
| S7 | 2.2 | 1.211 | 6.8 | 24.02 | 0.117 | 2.827 | 0.05 | 0.067 | 12.067 | 123.464 | 19.497 | 0.078 | 2.011 | 5.307 |
| S8 | 2.3 | 0.978 | 5.93 | 25.64 | 0.544 | 7.933 | 0.112 | 0.079 | 27.459 | 186.601 | 46.469 | 0.224 | 2.706 | 9.307 |
| S9 | 2.4 | 0.613 | 7.1 | 26.824 | 0.286 | 4.845 | 0.119 | 0.235 | 29.105 | 73.241 | 86.561 | 6.441 | 1.948 | 13.241 |
| S10 | 2.5 | 0.755 | 6.4 | 55.743 | 0.59 | 15.717 | 0.1 | 0.279 | 80.24 | 167.265 | 69.062 | 0.419 | 6.387 | 9.78 |
| S11 | 3.1 | 1.948 | 9.41 | 34.531 | 0.34 | 4.316 | 0.067 | 0.012 | 28.752 | 22.891 | 27.683 | 0.123 | 2.02 | 7.96 |
| S12 | 3.2 | 0.125 | 5.1 | 4.649 | 0.128 | 2.78 | - | - | - | - | - | - | - | - |
| S13 | 3.3 | 0.174 | 4.78 | 7.364 | 0.091 | 1.798 | 0.064 | 0.155 | 2.191 | 151.275 | 36.255 | 0.211 | 1.873 | 1.434 |
| S14 | 3.4 | 0.129 | 6.14 | 20.157 | 0.116 | 3.437 | 0.314 | 0.266 | 2.107 | 187.753 | 126.282 | 0.596 | 7.316 | 6.72 |
| S15 | 3.5 | 0.199 | 4.71 | 7.132 | 0.102 | 2.885 | 0.115 | 0.338 | 2.266 | 256.461 | 95.905 | 0.318 | 3.022 | 2.068 |
| S16 | 4.1 | 0.632 | 10.3 | 21.667 | 0.202 | 2.808 | 0.092 | 0.016 | 5.491 | 9.98 | 24.85 | 0.036 | 1.042 | 2.725 |
| S17 | 4.2 | 0.159 | 4.9 | 3.969 | 0.088 | 2.03 | 0 | 0.012 | 1.964 | 116.27 | 10.913 | 0 | 1.567 | 0.714 |
| S18 | 4.3 | 0.191 | 4.8 | 12.915 | 0.33 | 8.78 | 0.06 | 0.239 | 4.573 | 239.96 | 107.952 | 0.398 | 6.362 | 3.777 |
| S19 | 4.4 | 0.28 | 5.2 | 16.851 | 0.096 | 2.485 | 0.04 | 0.1 | 0.922 | 145.531 | 4.489 | 0.16 | 1.764 | 1.483 |
| S20 | 4.5 | 0.2 | 5.5 | 11.151 | 0.143 | 3.236 | 0.088 | 0.052 | 1.242 | 131.904 | 55.912 | 0.156 | 2.164 | 2.445 |
| S21 | 5.1 | 1.595 | 9.22 | 78.99 | 0.652 | 6.515 | - | - | - | - | - | - | - | - |
| S22 | 5.2 | 0.125 | 5.71 | 6.372 | 0.184 | 2.919 | 0.123 | 0.158 | 4.277 | 98.059 | 53.703 | 0.202 | 2.218 | 2.535 |
| S23 | 5.3 | 0.129 | 5.52 | 10.772 | 0.114 | 1.933 | 0.131 | 0.207 | 2.425 | 92.286 | 62.028 | 0.191 | 2.425 | 2.863 |
| S24 | 5.4 | 0.083 | 5.6 | 6.35 | 0.133 | 2.743 | 0.163 | 0.198 | 3.095 | 124.841 | 62.46 | 0.262 | 3.532 | 2.54 |
| S25 | 5.5 | 0.199 | 6.3 | 14.082 | 0.108 | 2.043 | 0.191 | 0.231 | 1.948 | 117.137 | 119.085 | 0.338 | 3.181 | 4.414 |
| S26 | 6.1 | 0.809 | 9 | 22.404 | 0.228 | 2.618 | 0.024 | 0 | 45.129 | 9.94 | 17.137 | 0.024 | 0.557 | 8.509 |
| S27 | 6.2 | 0.369 | 6.6 | 16.941 | 0.31 | 7.563 | 0.06 | 0 | 1.076 | 86.454 | 28.685 | 0.1 | 2.59 | 6.574 |
| S28 | 6.3 | 0.211 | 4.8 | 8.381 | 0.204 | 5.188 | 0.002 | 0.06 | 5.976 | 205.378 | 54.183 | 0.179 | 6.175 | 3.586 |
| S29 | 6.4 | 0.214 | 4.2 | 9.362 | 0.468 | 18.18 | 0.139 | 0.319 | 9.562 | 544.821 | 82.271 | 0.757 | 15.538 | 10.757 |
| S30 | 6.5 | 0.174 | 4.6 | 6.525 | 0.25 | 7.039 | 0.008 | 0.239 | 2.386 | 319.881 | 72.565 | 0.278 | 9.94 | 2.386 |
| S31 | 7.1 | 0.614 | 8.5 | 13.879 | 0.125 | 1.349 | 0.012 | 0.016 | 68.254 | 10.794 | 9.921 | 0.032 | 2.063 | 7.738 |
| S32 | 7.2 | 0.704 | 7.5 | 5.967 | 0.172 | 2.199 | 0.089 | 0.02 | 0.605 | 24.355 | 16.653 | 0.085 | 0.685 | 1.048 |
| S33 | 7.3 | 0.529 | 8.6 | 15.612 | 0.123 | 1.165 | 0.036 | 0.016 | 9.206 | 10.119 | 11.032 | 0.032 | 0.714 | 2.063 |
| S34 | 7.4 | 0.491 | 8.1 | 22.8 | 0.212 | 2.208 | 0.01 | 0 | 7.769 | 11.355 | 16.932 | 0.014 | 1.275 | 2.191 |
| S35 | 7.5 | 0.495 | 8.8 | 17.509 | 0.15 | 1.475 | 0 | 0 | 1.72 | 6.8 | 9.6 | 0 | 0.62 | 0.86 |
| S36 | 8.1 | 0.368 | 8.97 | 23.217 | 0.199 | 1.806 | 0.052 | 0.024 | 7.92 | 12 | 19.8 | 0.076 | 1.28 | 1.72 |
| S37 | 8.2 | 0.249 | 6.59 | 2.678 | 0.189 | 2.3 | 0.108 | 0.116 | 24.502 | 66.693 | 74.422 | 0.219 | 2.351 | 2.669 |
| S38 | 8.3 | 0.15 | 6.26 | 4.01 | 0.185 | 3.232 | 0.12 | 0.179 | 40.717 | 164.502 | 44.821 | 0.295 | 3.586 | 2.51 |
| S39 | 8.4 | 0.145 | 6.57 | 4.841 | 0.143 | 2.083 | 0.079 | 0.175 | 18.016 | 67.619 | 75.675 | 0.163 | 2.46 | 2.103 |
| S40 | 8.5 | 0.12 | 6.28 | 1.595 | 0.077 | 1.035 | 0.068 | 0.084 | 33.653 | 64.311 | 76.527 | 0.16 | 1.637 | 0.878 |
| S41 | 9.1 | 1.19 | 9.9 | 4.225 | 0.043 | 0.42 | 0 | 0 | 15.149 | 1.948 | 9.503 | 0 | 0.398 | 2.744 |
| S42 | 9.2 | 0.842 | 7.9 | 11.879 | 0.193 | 2.224 | 0.044 | 0.032 | 37.813 | 16.978 | 15.547 | 0.151 | 6.163 | 6.004 |
| S43 | 9.3 | 0.399 | 6.5 | 6.765 | 0.219 | 2.839 | 0.085 | 0.048 | 3.139 | 45.875 | 20.523 | 0.113 | 1.529 | 1.972 |
| S44 | 9.4 | 1.075 | 7.6 | 25.192 | 0.473 | 7.197 | 0.006 | 0 | 76.305 | 56.024 | 25.301 | 0.008 | 5.823 | 11.847 |
| S45 | 9.5 | 0.994 | 7.4 | 27.615 | 0.412 | 6.18 | 0.004 | 0 | 41.905 | 55.782 | 26.939 | 0.027 | 1.361 | 14.15 |
| S46 | 10.1 | 1.492 | 10.37 | 3.92 | 0.041 | 0.378 | 0.016 | 0.008 | 26.851 | 2.416 | 24 | 0 | 0.634 | 3.129 |
| S47 | 10.2 | 0.038 | 6.38 | 4.962 | 0.098 | 0.88 | 0.076 | 0.052 | 3.6 | 24.8 | 16.8 | 0.092 | 0.72 | 1.12 |
| S48 | 10.3 | 0.084 | 5.55 | 8.396 | 0.167 | 2.64 | 0.184 | 0.132 | 2.754 | 119.521 | 56.487 | 0.232 | 3.473 | 3.792 |
| S49 | 10.4 | 0.113 | 5.38 | 11.135 | 0.299 | 5.348 | 0.262 | 0.31 | 2.068 | 174.274 | 161.511 | 0.302 | 6.203 | 7.356 |
| S50 | 10.5 | 0.619 | 5.2 | 23.804 | - | - | 0.191 | 0.336 | 15.245 | 487.339 | 92.868 | 0.724 | 10.801 | 12.765 |
| S51 | 11.1 | 0.586 | 6.7 | 4.79 | 0.109 | 1.211 | 0.032 | 0.012 | 32.575 | 36.766 | 12.375 | 0.519 | 1.277 | 8.822 |
| S52 | 11.2 | 0.143 | 5.3 | 8.341 | 0.141 | 2.234 | 0.044 | 0.024 | 1.76 | 108.44 | 19.08 | 0.112 | 0.76 | 1.68 |
| S53 | 11.3 | 0.068 | 5.5 | 38.931 | 0.412 | 6.926 | 0.228 | 0.12 | 0.319 | 185.15 | 20.639 | 0.259 | 4.79 | 8.743 |
| S54 | 11.4 | 0.108 | 5.2 | 8.733 | 0.289 | 6.522 | 0.151 | 0.254 | 3.73 | 189.444 | 56.786 | 0.302 | 4.444 | 4.286 |
| S55 | 11.5 | 0.079 | 5.5 | 5.168 | 0.189 | 3.58 | 0.115 | 0.166 | 1.465 | 148.713 | 37.03 | 0.23 | 3.644 | 2.139 |
| S56 | 12.1 | 0.469 | 7.14 | 17.812 | 0.042 | 0.396 | 0.016 | 0.024 | 25.07 | 15.489 | 31.178 | 0.08 | 3.433 | 3.473 |
| S57 | 12.2 | 0.346 | 5.08 | 4.741 | 0.097 | 1.352 | 0.052 | 0.079 | 3.254 | 82.143 | 15.833 | 0.131 | 1.349 | 1.786 |
| S58 | 12.3 | 0.225 | 4.88 | 13.569 | 0.344 | 5.101 | 0.187 | 0.254 | 75.706 | 497.018 | 73.042 | 0.517 | 7.078 | 5.288 |
| S59 | 12.4 | 0.094 | 5.31 | 1.934 | 0.093 | 1.789 | 0.083 | 0.239 | 94.871 | 174.553 | 34.871 | 0.382 | 1.909 | 1.193 |
| S60 | 12.5 | 0.187 | 4.82 | 14.771 | 0.268 | 3.748 | 0.112 | 0.148 | 1.477 | 350.1 | 23.353 | 0.311 | 5.908 | 2.954 |
| S61 | 13.1 | 0.537 | 9.8 | 3.522 | 0.038 | 0.518 | 0.003 | 0 | 21.71 | 4.652 | 10.457 | 0.001 | 0.382 | 4.573 |
| S62 | 13.2 | 0.408 | 4.7 | 25.294 | 0.535 | 7.757 | 0.099 | 0.179 | 5.765 | 320.676 | 51.491 | 0.398 | 8.35 | 8.151 |
| S63 | 13.3 | 0.122 | 5.1 | 8.116 | 0.205 | 4.422 | 0.139 | 0.099 | 3.175 | 209.325 | 53.175 | 0.456 | 6.151 | 5.159 |
| S64 | 13.4 | 0.073 | 5.8 | 3.571 | 0.075 | 1.54 | 0 | 0 | 8.485 | 51.313 | 18.384 | 0.04 | 0.606 | 0.485 |
| S65 | 13.5 | 0.093 | 5.3 | 6.04 | 0.093 | 2.276 | 0.02 | 0.04 | 2.794 | 119.96 | 35.729 | 0.16 | 1.796 | 2.395 |
| S66 | 14.1 | 0.087 | 6.08 | 5.747 | 0.161 | 4.012 | 0.356 | 0.166 | 3.564 | 160.475 | 168.079 | 0.594 | 3.248 | 7.129 |
| S67 | 14.2 | 0.057 | 6.46 | 7.404 | 0.231 | 4.794 | 0.371 | 0.439 | 0.878 | 151.537 | 369.261 | 0.719 | 2.675 | 7.106 |
| S68 | 14.3 | 0.041 | 6.36 | 6.045 | 0.144 | 2.632 | 0.397 | 0.246 | 3.492 | 116.786 | 227.381 | 0.675 | 2.103 | 7.976 |
| S69 | 14.4 | 0.048 | 5.89 | 3.65 | 0.127 | 2.656 | 0.207 | 0.215 | 0.837 | 105.219 | 131.155 | 0.371 | 2.271 | 4.024 |
| S70 | 14.5 | 0.069 | 6.25 | 7.487 | 0.176 | 4.066 | 0.394 | 0.319 | 1.514 | 185.578 | 256.175 | 0.757 | 3.187 | 7.928 |

**Table S3** FUNGuild assignment breakdown of OTUs

| **Fungal Guild** | **# OTUs** |
| --- | --- |
| Algal Parasite--Leaf Saprotroph-Wood Saprotroph | 1 |
| Algal Parasite-Fungal Parasite-Undefined Saprotroph | 1 |
| Animal Endosymbiont-Animal Pathogen-Endophyte-Plant Pathogen-Undefined Saprotroph | 1 |
| Animal Pathogen | 2 |
| Animal Pathogen-Dung Saprotroph-Endophyte-Lichen Parasite-Plant Pathogen-Undefined Saprotroph | 6 |
| Animal Pathogen-Endophyte-Fungal Parasite-Plant Pathogen-Wood Saprotroph | 2 |
| Animal Pathogen-Endophyte-Lichen Parasite-Plant Pathogen-Soil Saprotroph-Wood Saprotroph | 3 |
| Animal Pathogen-Endophyte-Plant Pathogen-Undefined Saprotroph | 1 |
| Animal Pathogen-Endophyte-Plant Pathogen-Wood Saprotroph | 6 |
| Animal Pathogen-Fungal Parasite-Undefined Saprotroph | 8 |
| Animal Pathogen-Plant Pathogen-Undefined Saprotroph | 3 |
| Animal Pathogen-Undefined Saprotroph | 3 |
| Arbuscular Mycorrhizal | 6 |
| Bryophyte Parasite-Dung Saprotroph-Ectomycorrhizal-Fungal Parasite-Leaf Saprotroph-Plant Parasite-Undefined Saprotroph-Wood Saprotroph | 1 |
| Dung Saprotroph | 5 |
| Dung Saprotroph-Endophyte-Litter Saprotroph-Undefined Saprotroph | 3 |
| Dung Saprotroph-Plant Parasite-Soil Saprotroph-Undefined Saprotroph-Wood Saprotroph | 1 |
| Dung Saprotroph-Plant Saprotroph | 1 |
| Dung Saprotroph-Plant Saprotroph-Wood Saprotroph | 1 |
| Dung Saprotroph-Soil Saprotroph-Undefined Saprotroph | 5 |
| Ectomycorrhizal | 1 |
| Ectomycorrhizal-Endophyte-Ericoid Mycorrhizal-Litter Saprotroph-Orchid Mycorrhizal | 27 |
| Ectomycorrhizal-Fungal Parasite | 1 |
| Ectomycorrhizal-Fungal Parasite-Plant Pathogen-Wood Saprotroph | 3 |
| Ectomycorrhizal-Fungal Parasite-Soil Saprotroph-Undefined Saprotroph | 3 |
| Ectomycorrhizal-Lichenized-Wood Saprotroph | 1 |
| Endomycorrhizal-Plant Pathogen-Undefined Saprotroph | 3 |
| Endophyte | 2 |
| Endophyte-Litter Saprotroph-Soil Saprotroph-Undefined Saprotroph | 3 |
| Endophyte-Plant Pathogen | 1 |
| Endophyte-Plant Pathogen-Undefined Saprotroph | 2 |
| Endophyte-Plant Pathogen-Wood Saprotroph | 1 |
| Epiphyte-Litter Saprotroph | 2 |
| Ericoid Mycorrhizal | 5 |
| Fungal Parasite | 2 |
| Fungal Parasite-Plant Pathogen-Plant Saprotroph | 3 |
| Leaf Saprotroph-Plant Pathogen-Undefined Saprotroph-Wood Saprotroph | 7 |
| Lichenized | 3 |
| Orchid Mycorrhizal | 2 |
| Plant Pathogen | 17 |
| Plant Pathogen-Plant Saprotroph | 1 |
| Plant Pathogen-Undefined Saprotroph | 1 |
| Plant Saprotroph | 1 |
| Plant Saprotroph-Wood Saprotroph | 1 |
| Soil Saprotroph | 1 |
| Soil Saprotroph-Undefined Saprotroph | 1 |
| Undefined Saprotroph | 70 |
| Undefined Saprotroph-Undefined Biotroph | 1 |
| Undefined Saprotroph-Wood Saprotroph | 2 |
| Wood Saprotroph | 6 |
